# Brain-inspired spiking neural network controller for a neurorobotic whisker system

**DOI:** 10.1101/2021.10.26.465919

**Authors:** Alberto Antonietti, Alice Geminiani, Edoardo Negri, Egidio D’Angelo, Claudia Casellato, Alessandra Pedrocchi

**Affiliations:** Neurocomputational laboratory, Department of Brain and Behavioral Sciences, University of Pavia, Pavia, Italy; Nearlab, Department of Electronics, Information and Bioengineering, Politecnico di Milano, Milano, Italy; Brain Connectivity Center, IRCCS Mondino Foundation, Pavia, Italy

**Keywords:** Point neuron model, neurorobotic architecture, active whisking, Vibrissae, Trigeminal Ganglion, Trigeminal Nuclei, Facial Nuclei, CPG

## Abstract

It is common for animals to use self-generated movements to actively sense the surrounding environment. For instance, rodents rhythmically move their whiskers to explore the space close to their body. The mouse whisker system has become a standard model to study active sensing and sensorimotor integration through feedback loops. In this work, we developed a bioinspired spiking neural network model of the sensorimotor peripheral whisker system, modelling trigeminal ganglion, trigeminal nuclei, facial nuclei, and central pattern generator neuronal populations. This network was embedded in a virtual mouse robot, exploiting the Neurorobotics Platform, a simulation platform offering a virtual environment to develop and test robots driven by brain-inspired controllers. Eventually, the peripheral whisker system was properly connected to an adaptive cerebellar network controller. The whole system was able to drive active whisking with learning capability, matching neural correlates of behaviour experimentally recorded in mice.

## Introduction

A fundamental question in system neuroscience is to identify how peripheral sensory stimuli are processed in multiple brain regions showing specific neuronal activity. Rodent whisker-mediated touch system is a structurally well-known system which gives rise to complex adaptive behaviours (1). Specifically, the rodent whisker system represents an efficient combination of active perception and sensorimotor integration, in which self-generated movements are used to actively sense their environment, i.e., scanning the surrounding to collect behaviourally-relevant information. Rodents have specialized muscles in their mystacial pad able to control the hair position (2). They rhythmically protract their whiskers swiping the space surrounding the head and gathering information about shape and position of objects around them.

In the rodent whisker system, the primary afferences come from the trigeminal ganglion (TG) and the efferences project to motoneurons in the facial nuclei (FN). There are no direct connections between the two, indeed the innermost feedback loop is a di-synaptic reflex at brainstem level, involving interneurons from the trigeminal nuclear complex (TN) (3, 4). Outer loops in the whisker system involve multiple brain regions: the cerebellum, the midbrain (superior colliculus), and forebrain (5). The most extensive and most studied cortical loop is the whisker-barrel loop, involving the vibrissa primary sensory and motor cortex (6). Therefore, this somatosensory system is an ideal candidate to investigate the link between circuitry and function and to understand the underlying neuronal mechanisms in sensory readout and information processing.

Computational models of this system can play a fundamental role for multi-scale investigations, from neuron to behaviour, also thanks to availability of multi-scale experimental data in rodents for constraining and validating the models. In this work, we have developed a Spiking Neural Network (SNN) model able to process information encoded during whisking using a time coding representation of neuronal activity (7–10). While other models and kinds of artificial neural networks (e.g., rate-based or mean field models) are very powerful tools, based on brain dynamics, we choose SNNs because they are closer to biological reality, since they mimic the way information is coded and transmitted inside a real brain. In this work, each neuron in the network have been modeled with the most simplified spiking model, which is the Integrate & Fire model (I&F). SNNs can learn patterns of activity thanks to embedded plasticity models: here we included a Spike-Timing Dependent Plasticity (STDP) model (11–15) in the cerebellar circuit, which was inserted in the control system (outer loop) to test learning capabilities.

### Neurorobotic models of rodent whisking

Models of brain regions embedded in neurorobots allow to reproduce the functional mechanisms of living beings in closed perception-action loops (16, 17). Various examples of neurorobots using biologically inspired whiskers have been implemented in the last years. Among them, it is worth to cite the Whiskerbot, the SCRATCHbot and the Shrewbot (18).

The Whiskerbot consists of a robotic platform constituted by a head sensory unit of 150×170 mm and a two-wheeled body. The head carries six whiskers per side arranged in rows of three. Analogue information from whisker deflection is converted in empirically-based spike trains. It is able to freely move in an environment, actively whisking and orient toward salient stimuli using a neural network model of the superior colliculus (18, 19). The SCRATCHbot has a larger number of whiskers and degrees of freedom to position them in the environment. It was developed to reproduce different models of whisking pattern generation and to actively explore its environment using a simple model of tactile attention (18, 20). Both these robots were further enriched integrating the Shrewbot platform, which introduced algorithms able to detect texture and object from an active whisker array (18, 21, 22).

Real neurorobots are excellent test benches to challenge a neuro-inspired controller to demonstrate its capabilities, especially because of the real time computation constraints and the noise of the physical hardware and equipment, both intrinsic (non-ideal electronics sensors, limited spatio-temporal resolution, delays) and extrinsic (unexpected changes in the environment, external perturbing forces/torques, etc.). However, the implementation of physical neurorobots is complex and expensive, therefore limiting their adoption by neuroscientists to test computational models of the brain circuitry. Besides, it is also difficult to replicate the obtained results without an exact replica of the equipment used. Finally, the brain-inspired circuit controlling the robot can have a limited complexity, in terms of realism (neuronal models), number of elements (neurons and synapses), activity (spike events), and functionality (e.g., short and long term plasticity rules) for the sake of limited computational load required for real-time computations.

In this paper, we have developed a biologically-inspired neurorobotic whisker system on a virtual mouse inside the Neurorobotics Platform (NRP) (23–26). This work focuses on the reproduction of the peripheral parts of the whisker sensorimotor system and on the integration of the sensory inputs with an adaptive cerebellar spiking controller to perform a spatial learning task.

## Materials and methods

In this section, anatomy and physiology of the rodent whisker system are described and for each peripheral component (active vibrissae, sensory pathway, motor pathway, and trigeminal loop) the neurorobotic implementation is reported. Then the protocol to test the whisking controller, including an adaptive cerebellar network, is described, tailored on the experimental paradigms used on mice to understand neural mechanisms of active whisking and reward-based learning. Finally, the software libraries and computing resources are reported.

### Rodent whisker system and its neurorobotic implementation

Given the low number of degrees of freedom involved and the ease to make tests in laboratory conditions, the rodents whisker system has become a popular model for studying brain development, experience-dependent plasticity, active sensation, motor control, and sensorimotor integration (2, 5).

We have implemented the physical and the neural elements that constitute the whisker system of a rodent. The first step was the implementation of active whiskers (or vibrissae) in the mouse robot, making them controllable and allowing the reading of dynamic and kinematic parameters and information about the contact with external objects. Then, it was necessary to read inputs from the simulated environment and encode them realistically in the behaviour of vibrissal afferents. Once unprocessed data were gathered from afferents, further elaboration steps were carried out. These processed signals were used to directly control the motor actions, thus closing the first sensorimotor feedback loop, or extracting higher level information such as the phase of the whisking when a contact happened.

#### Active vibrissae

Vibrissae are long and sensitive hairs common to most mammals, including all primates except humans (27). Mystacial vibrissae are the ones growing on the mystacial pad, located at the sides of the animal snout, and have a major role in tactile spacial sensing and object discrimination (28).

### Neurorobotic implementation

To implement sensible whiskers in the mouse robot model inside the Neurorobotic Platform, we started from the HBP Mouse Robot v2 (dimensions: 140 cm from the nose tip to the end of the tail, 35 cm width, 35 cm height). The 3D models of the whiskers were defined as rigid cylinders: two right and two left whiskers, anchored to the mouse nose and with two different lengths and roll angles (Figure 1A). The lower whiskers (L0 and R0 for left and right whisker, respectively) are longer (50 cm each) and are rotated of 11°, while the upper whiskers (L1 and R1) are shorter (25 cm each) and are rotated of 22°. All whiskers have a diameter of 1 cm and their position can be independently controlled setting a torque at the revolving joints that link them to the mouse nose.

**Fig. 1.**
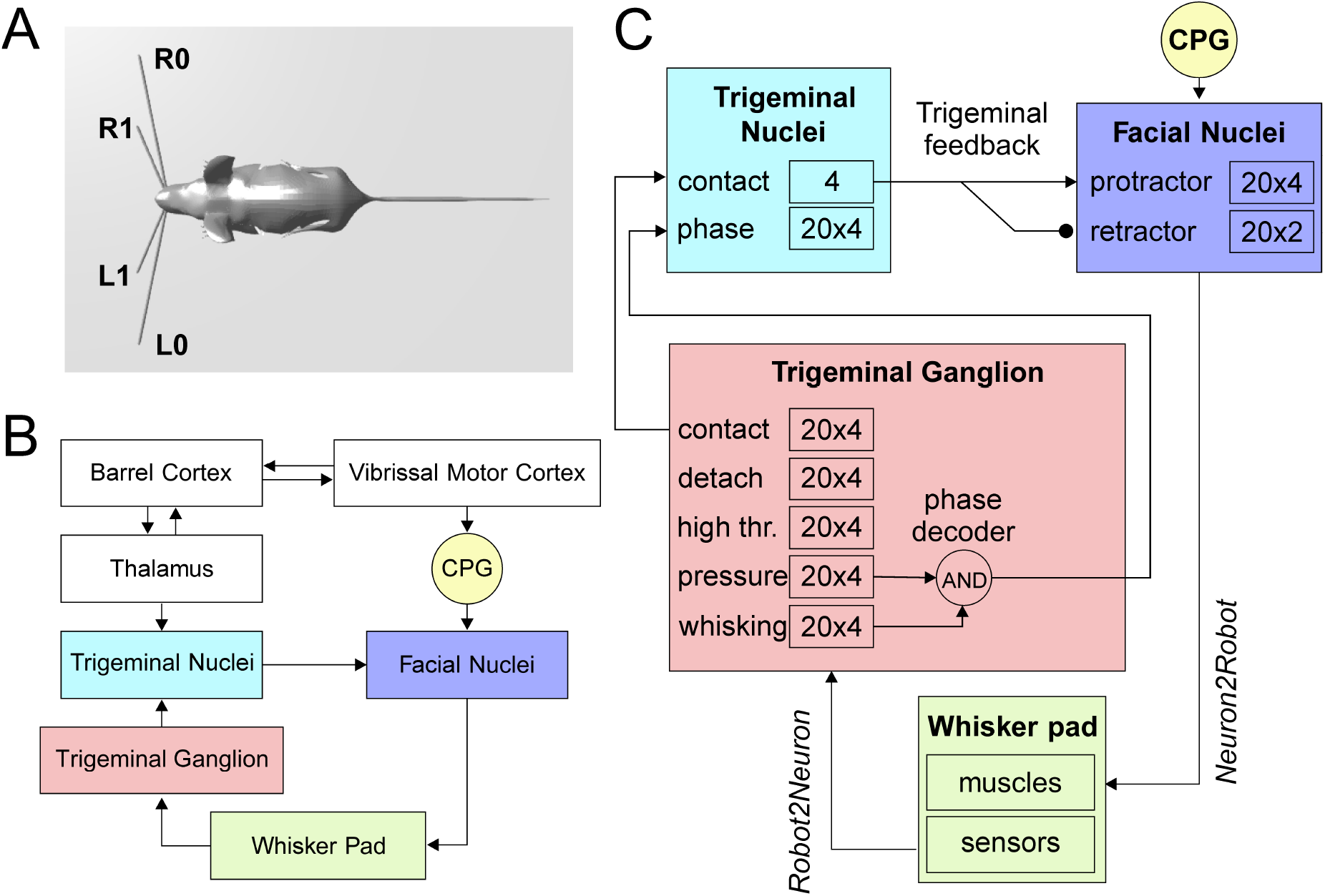
The rodent whisker system. A) Virtual robotic mouse implemented in the NRP, with two whiskers per side. L0 and R0 are the lower left and right whiskers, L1 and R1 are the upper whiskers. B) Block diagram of the rodent whisker system, including sensory and motor pathways, and its integration with higher-order areas (thalamus and cortex). C) SNN implementation of the mouse peripheral whisker system; numbers in each block represent the size of the neural populations included in that brain region. Arrows represent excitatory connections, circles inhibitory connections.

#### Sensory pathway

In the rodent whisker system (Figure 1B), the primary afferents have their nucleus located in the trigeminal ganglion.

When an object enters the peri-personal space around the head of a rodent, it will be sensed by the moving whiskers and its position and shape are encoded by neuronal signals. This can be done directly by the primary afferents or after some processing inside inner brain structures. The way the position of an external object is represented inside the vibrissal system of rodents is called vibrissal location coding (29).

The spatial organization of whiskers in the mystacial pad varies between different mammals, but are quite similar between rats and mice (28). Rats and mice whiskers are aligned in 5 rows, the upper 2 have four whiskers each, while the lower 3 counts 7 whiskers each (5). From rostral to caudal in each vibrissal row, the whisker length increases exponentially (28).

In the vibrissal follicle, three types of mechanoreceptors are present: Merkel cells, lanceolate endings and free nerve endings (5). Merkel cells ending are slowing adapting mechanoreceptors and they mostly signal ongoing movements, while lanceolate endings are rapidly adapting and respond to fast changes.

Mechanoreceptors surrounding each whisker transmit sensory information through cells whose bodies are located in the TG. Each neuron sends signals from a single vibrissa, while each follicle is innervated by ∼200 TG neurons.

A main distinction between trigeminal ganglion cells can be drawn based on the kind of receptor they receive information from: rapidly adapting or slowly adapting. While measuring cell activities *in-vivo* during free whisking, it can be observed that slowly adapting cells fire more during whisking in air. In fact, when contacting an object, all cells increase their firing rate, but the increase is more consistent for rapidly adapting cells (30).

In 2003, Szwed et al. induced artificial whisking in rats and measured the activity of TG cells (31). According to their results, TG neurons can be classified into distinct categories based on their responses to whisking in air and against an object:

- **Touch cells** responding only when the whisker touches an object, they can be further divided into **contact cells**, responding only at the beginning of the contact, **detach cells**, only at the end of the contact, and **pressure cells** with a tonic force-dependent response.
- **Whisking cells** responding only on whisker movement and not on object contact (if the contact does not affect the movement of the follicle).
- **High threshold cells** responding only to strong mechanical stimulations.

### Neurorobotic implementation

The Neurorobotics Platform permits to connect the environment to the robot sensors and actuators using so called transfer functions. These are Python functions defining to and from which ROS topics and neural populations read and write. They can be defined as *Robot*2*Neuron* (sensory) or *Neuron*2*Robot* (motor) according to the direction of the information flow.

The transfer function written to implement the follicle sensors is of the *Robot*2*Neuron* kind, since it reads the information on whisker mechanical status and position, and then process the data to obtain: possible contacts of a whisker against an object, the contact distance from the snout, the whisker angular position. The input transfer function is connected to the trigeminal ganglion neurons (Figure 1C), divided in five populations (TG pressure, TG high threshold, TG contact, TG detach, and TG whisking cells) with 20 neurons per whisker each, based on the functional separation observed in literature (31).

TG high threshold cells fire with a fixed rate when the contact is very close to the snout (less then 2 cm), constituting *defacto* a labelled line encoding for proximity, and TG whisking cells encode the current whisker position, each neuron has a Gaussian-shaped sensitivity and fire when the whisker position is within a narrow range around its maximum sensitive angle.

#### Motor pathway

The head of a rodent exploring its peripersonal space is constantly moving, side-to-side and up-and-down, while its nose moves side-to-side and the whiskers scan back-and-forth. These movements have a rhythmic component phase locked to sniffing, centered around 7 Hz in rats and 11 Hz in mice (6).

The representation of vibrissae in the primary motor cortex occupies around 20% of motor cortical area. There is no accepted topographic map, some studies obtained single-whisker responses, others observed how the number of whiskers showing evoked movements changes with the level of anesthesia used. *In-vivo* single cell microstimulation consistently evoked multi-whisker movements. There is strong evidence that the primary motor cortex controls only indirectly the muscle activity projecting to brainstem premotor networks, acting as central pattern generators (CPG) (32).

Motor neurons controlling muscles of the whisker pad are located in the lateral FN and send motor commands via the facial nerve. About 80% of the FN neurons evoking whisker movements induce protractions of a single whisker and about 20% the retraction of multiple whiskers (5).

### Neurorobotic implementation

The CPG has been implemented in the robot mouse as a single neuron. Controlled by a *Robot*2*N euron* transfer function, the CPG neuron emits regular spikes at a constant frequency in lower-theta band (4 *Hz*). It is connected with excitatory synapses to both protractors and retractors neurons, with delays of 1 *ms* and 50 *ms* respectively, in order to generate a rhythmic whisking movement.

Facial nuclei (Figure 1C) are divided into protractors and retractors. Protractors have been implemented with four populations of 20 neurons, where each population controls one whisker (L0, L1, R0, and R1). For retractors there are just two populations, one for each side (one population for L0 and L1, and one population R0 and R1). Sizes of populations are based on biological evidences, as described before.

The spiking activity of protractors and retractors is then transformed into a torque signal, applied to each whisker, using (1).

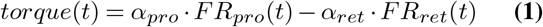

Where *FR*_*pro*_(*t*) and *FR*_*ret*_(*t*) are the instantaneous firing rates of protractors and retractors, respectively (in Hz), while *α*_*pro*_ and *α*_*ret*_ are constant gains, set to 0.15 Nm/Hz and 0.10 Nm/Hz, respectively.

#### Trigeminal loop

The trigeminal loop in the brainstem is a second order loop and is the most peripheral of the various loops constituting the vibrissal sensorimotor system. On the afferent side, neurons in the TG gather information from the follicles and project with excitatory synapses to the trigeminal nuclei complex. On the efferent side, subcortical whisking centers and CPG send motor commands to the motoneurons in the FN (3). Facial motoneurons driving muscles to protract the vibrissae receive a short latency input (7.5 ± 0.4ms) followed by synaptic excitation from neurons in TN. These connections result in a pull-push mechanism allowing for rapid modulation of vibrissa touch during exploration.

### Neurorobotic implementation

When a whisker touches an object, the physical simulator makes it bounce according to physical properties of the simulated materials, producing a noisy contact signal. This offers us the possibility to apply the trigeminal feedback mechanisms previously described as a biologically inspired debouncing mechanism. First, a neural population (TN contact) has been created in the trigeminal nuclei, of the same size of TG contact and taking from it excitatory, one-to-one connections (Figure 1C). Then, TN contact neurons were connected to the facial nuclei protractors with excitatory synapses having a 7.5 *ms* delay (4), and with inhibitory synapses to the retractors. This increases the joint torque sent to the colliding whiskers, in order to impede their rebound and to keep them at contact with the touched object.

In the TN, we have included an additional population made of 20 neurons for each whisker, that has to encode the phase of the whisking period at which the contact occurs. TG has been implemented as an array of coincidence detectors (phase decoder), one for each TG whisking neuron, gating them in a logical AND with TG pressure neurons. The result is a labelled-line encoding of contact phase. The phase decoder is implemented by a transfer function that takes input from afferents in TG (pressure and whisking cells) and projects to the TN phase population in the trigeminal nuclei (Figure 1C). For each TG whisking cell, the same spike rate is propagated downward only if pressure cells are firing. The phase information is needed for the precise localization, with respect to the mouse head, of the object touched by the whiskers.

Table 1 summarizes the connectivity between the different populations of the whisker system.

**Table 1.**
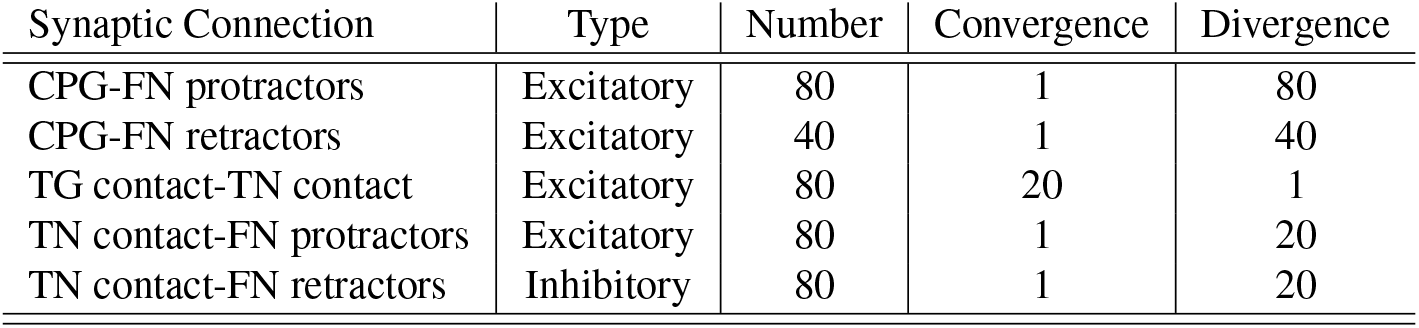
Connectivity of the SNN whisker system model

| Synaptic Connection | Type | Number | Convergence | Divergence |
| --- | --- | --- | --- | --- |
| CPG-FN protractors | Excitatory | 80 | 1 | 80 |
| CPG-FN retractors | Excitatory | 40 | 1 | 40 |
| TG contact-TN contact | Excitatory | 80 | 20 | 1 |
| TN contact-FN protractors | Excitatory | 80 | 1 | 20 |
| TN contact-FN retractors | Inhibitory | 80 | 1 | 20 |

#### Testing protocol

The peripheral components of the whisker system described above has been tested inside the Neurorobotics Platform in free whisking conditions. The mouse moved its whiskers in an empty environment or touching an object and the spiking activity of FN, TG and TN has been recorded in order to verify the proper functioning of the developed system.

##### Experimental whisking-based object localization task

To provide a meaningful example of how the developed mouse whisker system can be used to build *in-silico* neurorobotic experiments, we reproduced an experimental study which investigated the involvement of the cerebellum in a whisking-based object localization task in head-fixed mice (33). Rahmati and colleagues tested two populations of mice, one wild-type (Control) and one knock-out (L7-PP2B), suffering from genetically impaired cerebellar plasticity. Water-deprived mice had to learn to locate a vertical bar in their whisker field and lick a water droplet (GO trial) within a time response window or to refrain from licking (NOGO trial) according to the bar position.

During the first sessions, both mouse populations started with high hit rates and high false alarm rates. Control mice showed faster learning capabilities, reducing their licking response to NOGO trials after the first 4 training sessions. Conversely, knock-out mice reduced their licking in a random fashion staying close to the guess rate (33). Therefore, they concluded that cerebellar plasticity has a crucial role in this sophisticated cognitive task requiring strict temporal processing.

##### Cerebellar SNN model

To investigate *in-silico* the role of cerebellar plasticity during the task, we integrated a well established cerebellar-inspired SNN model to the whisker system described above. Recently, a detailed spiking neural network model of the cerebellar microcircuit proved able to reproduce multiple cerebellar-driven tasks (26, 34–37). Here, we used the model to drive learning in the *in-silico* whisking-based object localization task.

The SNN cerebellar microcircuit (Figure 2) was populated with leaky Integrate&Fire neurons, distinguishing between different neural groups. Mossy Fibers (MFs), the input to the cerebellar module, encode the state of the body-environment system: the whisker current position and the localization of an eventual object, e.g., the cue signalling a GO trial. Therefore, MFs receive excitatory connections from TG pressure cells and TN phase cells. Granular Cells (GrCs) represent in a sparse way the input from the MFs. Inferior Olive neurons (IOs), the other input to the cerebellar module, encode the reward, which is provided when a response is correctly generated (i.e., in a GO trial). In fact, this neural population responds to attention or surprise signals. Purkinje Cells (PCs) integrate the sparse information coming from the GrCs through the Parallel Fibers (PFs), while Deep Cerebellar Nuclei (DCN), the only output of the cerebellar module, generate the response (i.e., the equivalent of licking). The firing rate of DCN is monitored and a response is detected when the firing rate exceeds a pre-defined threshold (i.e., 80 Hz). The network structure and connectivity are reported in Figure 2 and Table 2.

**Table 2.** Connectivity of the SNN cerebellar model

| Synaptic Connection | Type | Number | Convergence | Divergence |
| --- | --- | --- | --- | --- |
| TG pressure-MFs | Excitatory | 80 | 1 | 1 |
| TN phase-MFs | Excitatory | 80 | 1 | 1 |
| MFs-GrCs | Excitatory | 8000 | 4 | 80 |
| PFs-PCs | Excitatory | 115200 | 1600 | 58 |
| IOs-PCs | Teaching | 72 | 1 | 1 |
| MFs-DCN | Excitatory | 3600 | 100 | 36 |
| PCs-DCN | Inhibitory | 72 | 2 | 1 |

**Fig. 2.**
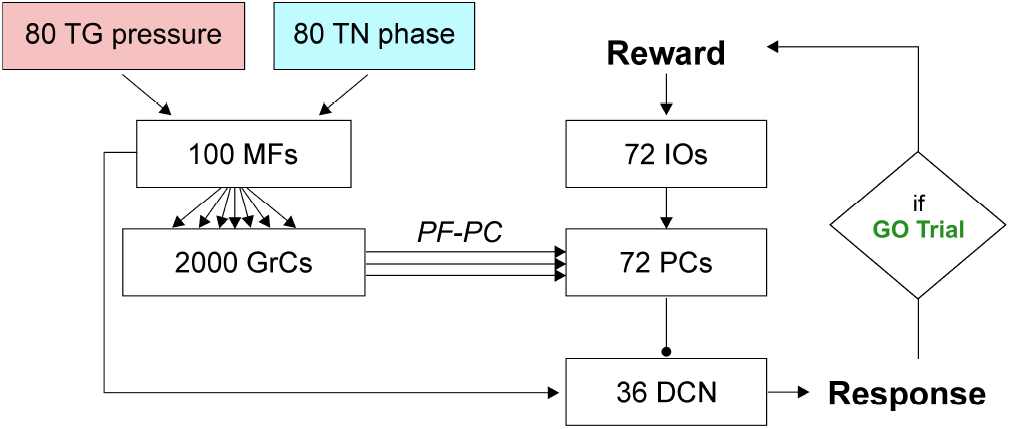
SNN implementation of the cerebellum. Whisking sensory signals are conveyed to the cerebellar MFs from TG pressure and TN phase neurons, while the reward signal during correct GO trials reach the IO neurons; the cerebellum controls the output motor response (head movement) according to DCN activity (i.e., generation of a response, head raise, when the firing rate exceeds a set threshold). Arrows and circles represent excitatory and inhibitory connections, respectively.

The cerebellar SNN model included one plasticity site, at the cortical level, between PFs and PCs, based on a well-known kind of STDP (38–40). Synaptic weights between PF-PC plasticity are modulated by IO activity (IOs-PCs connections in Table 2 are indicated as “teaching”), depending on the difference between the pre- and post-synaptic firing times (36, 41, 42). Long-Term Potentiation (LTP) and Long-Term Depression (LTD) are the two possible changes that each synaptic connection can undergo. Synaptic weights increase (LTP) whenever a PC only receives an input from a PF, while they decrease (LTD) when associated with IO inputs (43–47). The learning rule can be formalized as in (2).

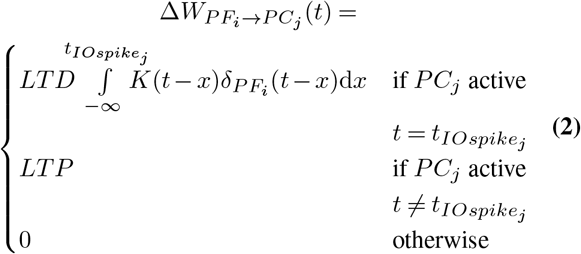

where:

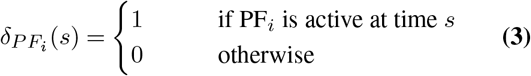

and the kernel function is:

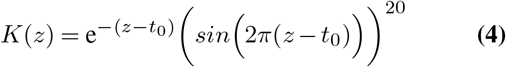

where 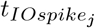 is the time when *IO*_*j*_ emits a spike; *K*(*z*) is the integral kernel function, which has its peak at *t*_0_ (100 ms) before 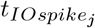. The plastic learning rule is characterized by two constants, *LT P* and *LT D*, which regulate the amount of synaptic change. These constants cannot be directly computed from physiological data, but they have been set to values found in related modelling studies (*LT P* = 0.01, *LT D* = −0.03) (37).

##### Neurorobotic implementation of the whisking-based object localization task

In the NRP, a licking-like movement, has been set for the virtual mouse: it has to rise its head and touch a shelf positioned just above. During GO trials, a vertical bar is displaced in the left whisker field and if the mouse rises its head, a reward signal is triggered. During NOGO trials the bar is on the right and if the mouse rises its head it does not receive any reward (Figure 3A).

**Fig. 3.**
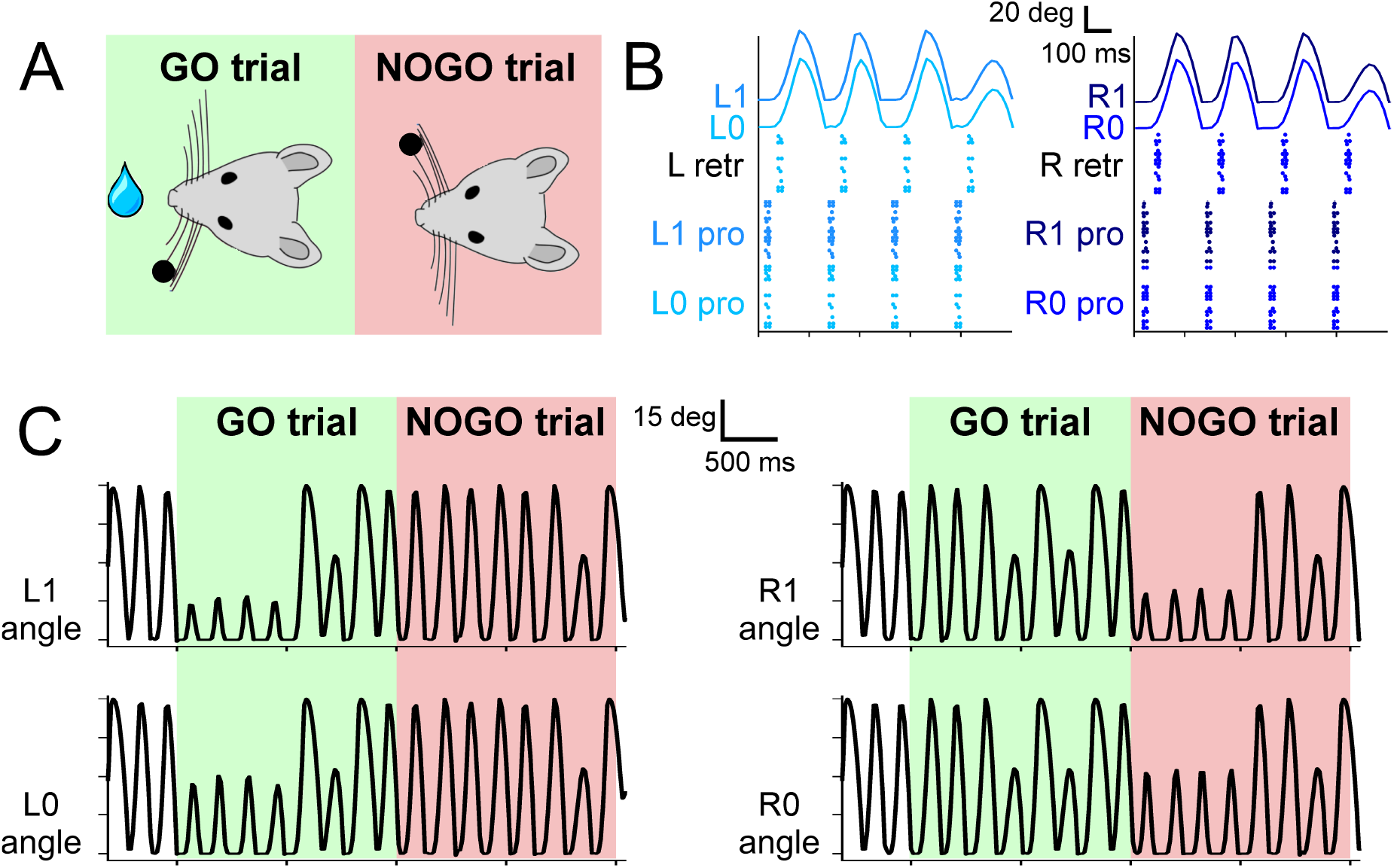
A) experimental protocol, during GO trials, a sensory cue (a small bar, depicted as a black dot) is placed in the left whisker field of the mouse. Correct responses lead to a reward (water drop). During NOGO trials, the sensory cue is placed in the right whisker field and a response does not result in any reward. B) Spiking activity of motor neurons for protraction and retraction during one trial. 20 protractors neurons for each whisker and 20 retractor neurons for each side (L and R) fire under the control of the CPG neuron at 4 Hz. The resulting displacement of each whisker is depicted in the upper part of the panel. C) Angular displacement of the four whiskers during one GO and one NOGO trial. During the GO trial, in the first second, left whiskers hit the sensory cue bar, placed in the left whisker field. On the other hand, during the NOGO trial, the right whiskers hit the sensory cue bar placed in the right whisker field.

The experiment is composed of short trials of 2 seconds, divided in GO and NOGO trials. The vertical bar is displayed in the mouse whisker field for 1 second, while the response window continue till the end of the trial.

Trials were grouped in sessions composed of 10 trials, 5 GO and 5 NOGO, performed in a randomized sequence. The neurorobotic experiment included 27 sessions, following the experimental protocol. In order to evaluate the learning of the controller, for each session we recorded the percentage of correct responses in GO trials (“hit rate”), and the percentage of responses in NOGO trials (“false alarms”).

##### Experimental and in-silico cerebellar impairment

PC-specific PP2B knock-out (L7-PP2B) mice show deficits in motor learning, consolidation, and procedural learning (48, 49) while they behave normally in standard non-motor tasks (50). In their experiment, Rahmati and colleagues tested how an impairment of the PF-PC LTP influenced the performance in the whisking-based object localization task. They demonstrated that learning in L7-PP2B mice was severely impaired, indicating that this task can depend, at least to some extent, on cerebellar plasticity.

We recreated *in-silico* the impaired cerebellum dramatically reducing the constant *LT P* (see (2)) to 10% (*LT P*_*L*7*−PP*2*B*_ = 0.001). We repeated each experiment (i.e., 27 sessions, 10 trials each, therefore 270 trials) of the localization protocol 10 times, using the impaired cerebellar model, then we compared the curves of hit rate and false alarms between L7-PP2B and control mice.

#### Hardware and software

For the simulations, we have used a local installation of the NRP version 3.1, exploiting Python 3.8 (RRID:SCR_008394), Gazebo 11 (51), and ROS Noetic (52).

The simulation of the controller has been done with NEST, a software simulator for spiking neural networks (53–55). We used NEST 2.18 (56) (RRID:SCR_002963), interfaced through PyNN 0.9.5 (57) (RRID:SCR_002963).

All the simulations have been carried out on a Desktop PC provided with Intel Core i7-2600 CPU @ 3.40 GHz and 16 GB of RAM, running 64 bit Ubuntu 20.04.2 LTS.

## Results

We successfully developed a SNN model of the sensorimotor peripheral whisker system, modelling trigeminal ganglion, trigeminal nuclei, facial nuclei, and central pattern generator neuronal populations. This peripheral SNN was embedded in a virtual mouse robot, and it was properly connected to an adaptive cerebellar SNN. The whole system was able to drive active whisking with learning capability, matching neural correlates of behaviour experimentally recorded in mice.

### Motor pathway

The four whiskers are controlled by the motoneurons present in the FN. They are working under the control of a single CPG neuron, firing at 4 Hz, which rhythmically excites protractors and retractors neurons. Motoneurons spikes are then transformed into torques applied independently at each whisker. As shown in Figure 3B, during a free whisking period, the spiking pattern of the four groups of neurons is very similar, with a precise temporal alternation between protractors, causing the whisker to move forward, and retractors, pulling the whiskers back to the initial position. Whiskers’ movements are slightly shifted with respect to the spikes due to the delays introduced by the conversion between spikes and torques and by the mechanical inertia of the whiskers. The mean firing rate of protractors and retractors neurons are 3.80 Hz and 4.00 Hz, respectively, with a peak firing rate of 48.75 Hz and 55.00 Hz, respectively. Figure 3C shows how the whisker trajectory changes during GO and NOGO trials. Namely, in GO trials a bar is placed for 1 second in the left whisker field, therefore whiskers L0 and L1 hit it and their range of motion is therefore reduced to ∼15 degrees. The same behaviour can be observed during NOGO trials for whiskers R0 and R1.

### Sensory pathway

Neurons in the TG and TN compose the sensory pathway and the first stage of elaboration is done by TG neurons. Each group of TG shows specific activity patterns depending on its function (Figure 4A). TG whisking neurons follow the angular profile of each of the four whiskers; it is possible to notice the differences between GO and NOGO trials, where left and right whiskers change their spatial profile when hitting the bar in the first second of each trial. Their mean firing rate is 3.65 (± 1.93) Hz. TG contact neurons fire at ∼5 Hz when the whisker hits the bar, while TG detach neurons when the whisker is no longer in contact with the bar, because the bar has been removed or because the whisker has been retracted. TG pressure neurons are active for the whole duration of the contact between the whisker and the bar (27.00 ± 17.69 Hz). In the SNN model, we have included an additional population, TG high threshold neurons, which are activated when the contact happens close to the nose of the robot (less than 2 cm). However, in our protocol the bar is placed at a higher distance, therefore those neurons were never activated.

**Fig. 4.**
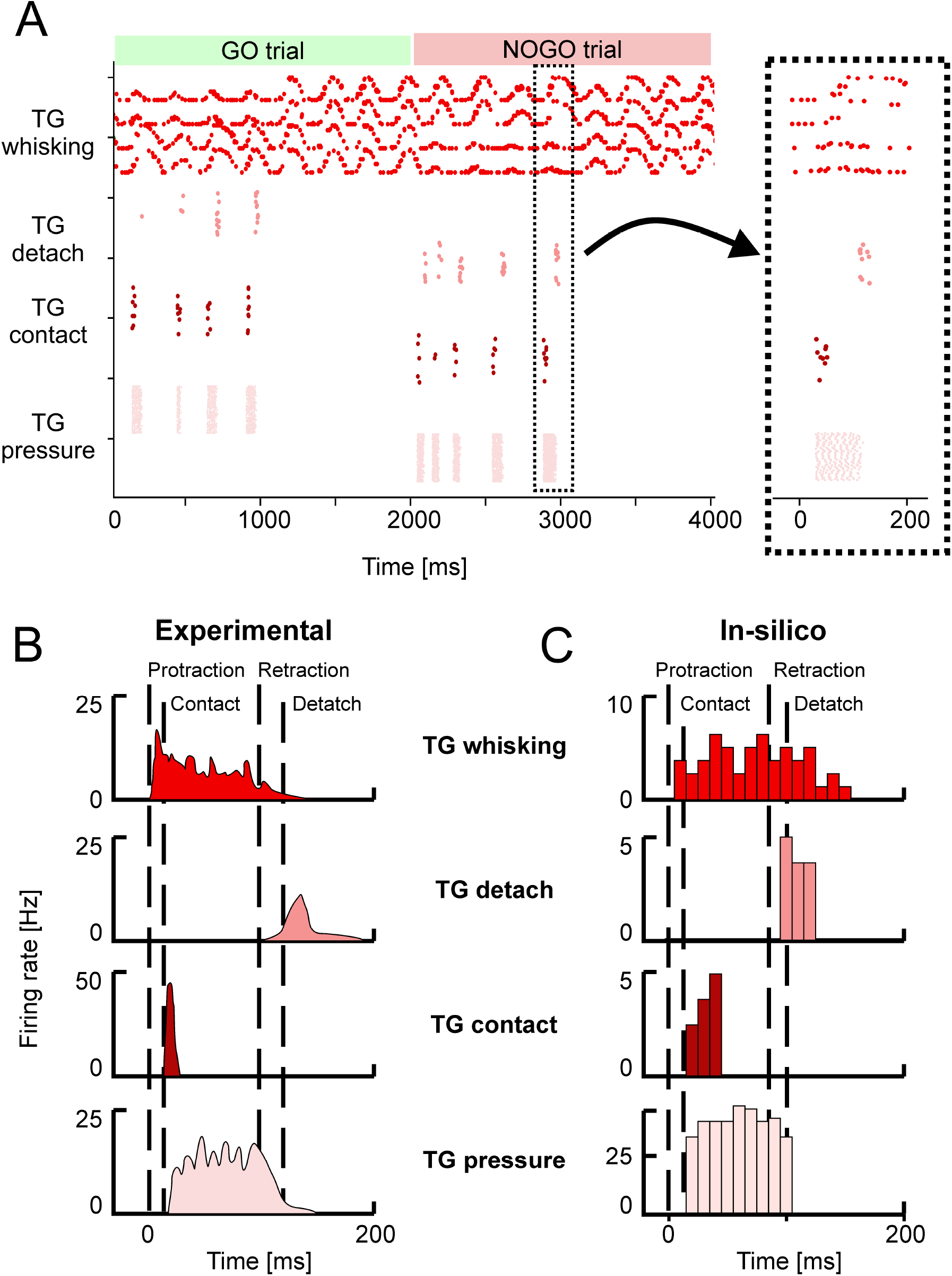
A) Spiking activity of the Trigeminal Ganglion (TG) neurons during GO and NOGO trials. Each row represents the activity of one neuron, different shades of red are used to plot the activity of the four groups of TG neurons. The inset shows a magnified portion of the full scatterplot, focusing on a single protraction-retraction movement of the whiskers. (B) Firing rates of TG populations measured experimentally during a single protraction-retraction movement, as reported in (29). Vertical dashed lines represent the four main events: start of the protraction, contact of the whisker against an object, start of the retraction, detach of the whisker from the object. (C) Firing rates recorded from the simulation of the SNN model of TG populations. The length of each bin is 10 ms. Colors are the same as panels A and B.

Figures 4B and 4C provide a direct comparison between the firing rates of the different TG populations during one whisker movement. Figure 4B has been adapted from (29), while in Figure 3C the firing rates of the neurons in a specific trial (the magnified inset from Figure 4A) have been computed with bins of 10 ms. It is possible to appreciate that *in-silico* TG neurons show a behaviour comparable to the one of biological neurons, especially for the timing of their response with respect to the events of protraction, contact, detach, and retraction.

### Learning performance

We have shown that the SNN representing the sensorimotor whisker system is able to encode in a biologically realistic way the sensory and motor signals exchanged with a robotic plant. To demonstrate how this system can be used to recreate a complex behavioural test, we connected the whisking sensory system to an adaptive SNN and we challenged the integrated system in the object localisation experiment proposed by Rahmati and colleagues (33). The aim of the mouse is to lick during the GO trials and to refrain from licking during the NOGO trials, distinguishing between the two conditions according to the position of a bar placed into their whisker field. Figure 5B reports the percentages of correct licks in GO trials and the number of incorrect licks in NOGO trials. The reference behavioural data recorded in animals are reported in Figure 5A.

**Fig. 5.**
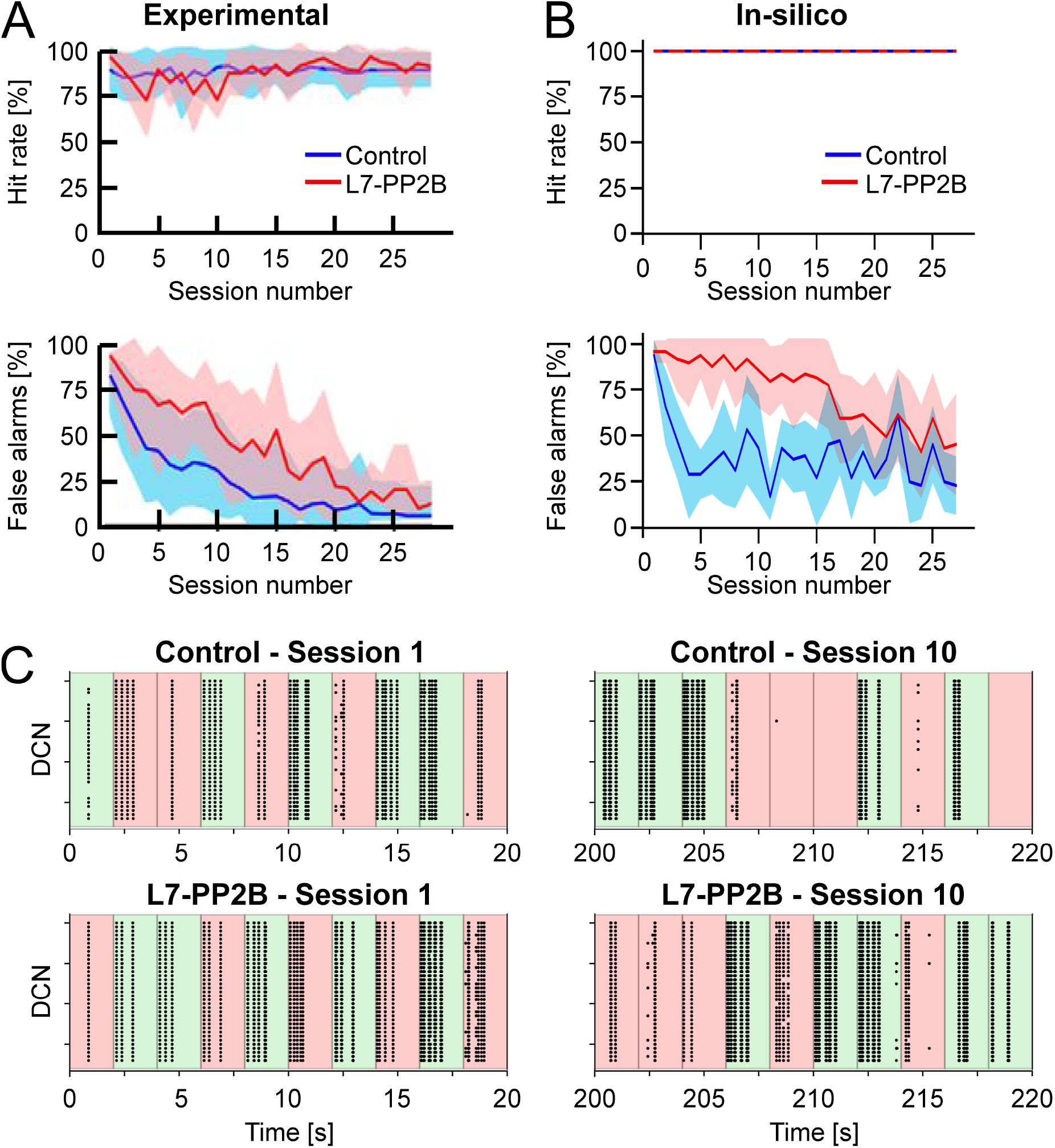
A) Learning curves recorded in the experiment performed by Rahmati and colleagues (33). The upper row shows the Hit rate (i.e., the percentage of correct responses in GO trials) along sessions, where each session is composed of 10 trials. The lower row shows the False alarms (i.e., the percentage of incorrect responses in NOGO trials) along sessions. The blue and red curves show the mean values for control animals and knock-out (L7-PP2B) mice. Shaded areas show the standard deviation. B) Learning curves recorded from the *in-silico* experiments (10 control and 10 knock-out models). Colors are the same as in panel A. C) Spiking activity of the DCN neurons during GO (green) and NOGO (red) trials, in the first session (left column) and after 10 sessions of training (right column). The first row reports the activity of one Control simulation, while the second row reports one knock-out simulation. Each dot is a spike of one of the 36 DCN in the cerebellar network. The order of GO and NOGO trials is randomized for each session and simulation, but all sessions have 5 GO and 50 NOGO trials.

Considering control animals, it is possible to see that mice lick continuously in the first sessions (i.e., hit rate and false alarms are both close to 100%), without distinguishing GO and NOGO trials. While the experiment proceeds, the animals learn to refrain from licking during the NOGO trials. In fact, the percentage of false alarms decreases toward 0%. At the same time, the percentage of correct licks remains close to 100%.

Control and L7-PP2B mice differed in their learning skills. Both started the training with high hit and false alarm rates. As a result, they performed close to the guess rate. During the subsequent sessions, control mice consistently increased accuracy, specifically reducing their response to NOGO trials. In contrast, L7-PP2B mice continue for more sessions to not discriminate between GO and NOGO trials. In the later sessions, also L7-PP2B diminished their licks in the NOGO trials, but their learning trajectories remained noisier than those of controls, taking a longer time to reach high performance levels. This observation suggested that a functional LTP mechanism was important to obtain a superior ability to rapidly discriminate between GO and NOGO cues, and then respond accordingly.

The performances and the learning trajectories of control and L7-PP2B *in-silico* models (Figure 5B) are similar to their biological counterparts. While both models maintain a perfect recognition of GO trials, the control model learned to refrain from licking in NOGO trial faster and stably. The variability present between the 10 different tests is due to the different sequences of GO and NOGO trials, which were randomly extracted for each session.

Looking at the spiking activity of the cerebellar network, and in particular at the DCN population (Figure 5C), which drives the response of the mouse, it is possible to appreciate the different evolution in the spiking patterns. Control and L7-PP2B simulations have a similar activity in the DCN during the 10 trials of the first session, the neurons fire regardless of the input arriving from the whisker system (right or left contacts) and therefore DCN generate a response during both GO and NOGO trials. After 10 sessions of training, the Control simulation shows intense DCN activity during GO trial and weak or null activity during NOGO trials, proving that the cerebellar network has learned the association between left/right stimulation with the presence/absence of the reward. This behaviour is impaired for the L7-PP2B simulation, in fact, there are still several DCN that are firing during both GO and NOGO trials, then causing a high False alarm rate.

Each session took about 140 minutes for a simulated time of 540 seconds (i.e., 270 trials of 2 seconds each), with a scale-up with respect to the real-time equal to ∼ 15 times. The maximum RAM consumption was equal to ∼ 15 GB.

## Discussion and conclusions

We developed a spiking neural model of the mouse whisker system, covering both sensory and motor pathways, and their interconnections. The implemented system takes into account the different roles that groups of cells have at the different stages of the sensorimotor processing, providing coding for complex information such as the object localization performed during the active whisking. This system, properly connected to an adaptive cerebellar-inspired spiking network, reproduced complex *in-vivo* experiments, by using the neuro-robotics platform.

The peripheral whisker system showed appropriate discharge patterns as in *in-vivo* experimental recordings during whisking, in precise time-windows of exploration and object interaction and depending on which side the stimulus was presented within the whisker field.

The peripheral system, when wired to a cerebellar SNN with plasticity and tested in an object-localization task, was able to reduce the number of useless responses along a sequence of trials (triggered by the NOGO trials) which did not correspond to any reward. This learning curve was slowed down when the plasticity parameter (LTP rate) of the cerebellar SNN was strongly reduced, as in knock-out mice recorded experimentally.

The integrated circuit, entirely made of spiking neurons, proved the good integration of different ways of neural coding. In fact, while the main parameter correlating response patterns to behavior was the average firing frequency of the DCN population, other elements of the whisker system used a variety of encoding strategies. For instance, the time-coded activity of TG contact and TG detach cells or the TG whisking cells encoding the current whisker position by means of their Gaussian-shaped sensitivity.

The model here proposed can be used as a reference for future advanced neurorobots and for neuroscience *in-silico* experiments, to investigate the role of cerebro-cerebellar loops and cerebellar physiology in whisking protocols.

### Limitations and future challenges

Considering the robotic aspects, a limitation of the physical simulator (Gazebo) regards the properties of the materials that can be used. Rodents rely on whiskers bending to recognise the shape of objects and on their resonance frequencies to detect textures (58, 59), but the current state of the simulators used by the NRP supports only rigid bodies. Using only rigid whiskers makes object recognition tasks more difficult, unless maybe using large arrays of finely spaced whiskers. Therefore, this work focused on extracting only spatial information, which can be easily performed with just rigid whiskers, and not more sophisticated features of the touched object.

Much of the work on the whisker system consisted in the encoding of information in TG primary afferents, ignoring all of the internal brain structures involved in the whisker system, in particular the somatotopic mapping emerging in the TN and propagated in the thalamo-cortical system. Loops between thalamus and cortex have been cited as possible location for mechanisms decoding phase information, with the use of neuronal phase-locking loops. A possible future development can be exploring other loops in the brainstem outside the TN, such as the ones involving the superior colliculus and their interactions with attention and foveation (60).

The work on the cerebellar control mechanisms was mainly limited by the long simulation times of the NRP, which influenced the choice of network and learning parameters. Given the limited number of mossy fibers and granule cells (100 and 2000, respectively), the cerebellar network showed a reduced generalization capability. The discrimination task, in this case, was between two very different conditions (object hit with the left or with the right whiskers). Mice have a higher resolution, since they can recognize objects slightly moved or even objects with different textures. With larger populations, training could make different sub-populations respond to different inputs, encoding for more complex features of the sensed environment. Future work can explore this hypothesis making rigorous analyses on cell responses and optimizing the network size to the variety of input patterns.

We have chosen I&F neuron models as building blocks of the SNN network. However, nowadays, there are much more complex models, taking into account many mechanisms related to membrane potential and ionic currents (biophysical models (61)) or more advanced I&F neuron models (62, 63). Even though these models are more accurate representations of the biological elements, their complexity would require too much computational power to simulate a network made of thousands of neurons, which would eventually prevent from embedding these models as controllers in neurorobots. Therefore, the model we are proposing is a trade-off between computational efficiency and biological realism.

NEST-based simulations offer a great possibility to develop and test biologically inspired models, but require high-performance computing for large-scale models (64) and therefore does not allow performances sufficient to control robots in real world situations. This could allow testing the robustness of the bioinspired controller against common environmental noise, increasing the similarity with experimental results (e.g., the imperfect (< 100%) hit rates accuracies achieved by experimental animals). An already available solution to achieve real-time performances can be to rely on spiking neural networks running on neuromorphic hardware. Very recently, a cerebellar-inspired model made of 97 thousand neurons and 4.2 million synapses has been implemented on the neuromorphic platform SpiNNaker (65). This type of solution could be applicable if the plasticity rule used at PFPC synapses, supervised by IO activity, will be implemented on this or other neuromorphic systems. On the other hand, the SNN whisker system here presented can be simulated on SpiNNaker chips, since it has been developed using PyNN, which supports both NEST and SpiNNaker as simulators.

### Conclusions

Neuroscientists have not fully uncovered the neural mechanisms for mouse whisking, but it is clear that it involves a complex architecture composed of multiple sensorimotor loops. In this work, we developed and tested a spiking computational model of the peripheral whisker system, reproducing the neural dynamics observed in its different components and embedded in a virtual mouse neurorobot controlled by a cerebellar SNN.

The virtual mouse enriched with this peripheral whisker system may be connected to more realistic multi-area brain models, to shed light on how these regions together may control the precise timing of whisker movements and coordinate whisker perception.

In the future, refined versions of the model could exhibit more advanced features, such as the recognition of surface textures, identification of movements of the touched object, or other complex touch-guided behaviours. From a technological perspective, neuromorphic implementations can be employed to speed up the computation until reaching real-time performances, granting the possibility to embed the whisker system in physical robots.

## Author Contributions

AA, AG, CC designed the model and the computational framework. AA, AG, EN carried out the implementation. AA and EN wrote the manuscript with input from all authors. AA, EDA, CC, AP conceived the study and were in charge of overall direction and planning. All authors discussed the results and commented on the manuscript.

## Data Availability Statement

The computational models used for the simulation are publicly available at https://github.com/alberto-antonietti/paper_whisking. Data generated from the simulations and Python codes to generate all the figures presented in this work are available in the same GitHub repository.

